# cMYC-mediated immune repression is reversed by inhibition of H3K9/H3K27 methylation maintenance

**DOI:** 10.1101/2023.10.18.562888

**Authors:** Isabel Dye, Sarah Laing, Ian Garner, Hasan B. Mirza, Nayana Iyer, Nicola Brady, Pavlina Spiliopoulou, Sarah Spear, James Robinson, Francesca Fiorentino, Matthew J. Fuchter, Daniel J. Murphy, Iain A. McNeish, Robert Brown

**Affiliations:** Dept Surgery & Cancer, Imperial College London, London W12 0NN, UK; School of Cancer Sciences, University of Glasgow, Glasgow, G61 1BD, UK; AstraZeneca, Cambridge Biomedical Campus, Cambridge CB2 0AA, UK; Nightingale-Saunders Clinical Trials & Epidemiology Unit (King’s Clinical Trial Unit), King’s College London, UK; Dept Chemistry, Imperial College London, London, W12 0NN, UK

## Abstract

Aberrant cMYC activity is a key driver of cancer, involved in several hallmark processes. Alongside the canonical hallmark of proliferation, cMYC represses immune signalling in a cell-intrinsic manner. The histone methyltransferases EZH2 and G9a interact with cMYC to modulate gene expression, including repression of immune genes via H3K27 and H3K9 histone methylation. Analyses of 565 cell lines derived from solid cancers demonstrated that greater cMYC-G9a/EZH2-mediated repression correlates with lower immune gene scores in a cell-intrinsic manner (innate, Type I and Type II IFN response), an effect most evident in *MYC*-amplified cell lines. In ovarian high-grade serous carcinoma (HGSC) cell lines and an *in vivo* murine model of HGSC, HKMTi-1-005, an inhibitor of H3K27/H3K9 methylation maintenance, relieved cMYC-G9a/EZH2 repression whilst inducing an immune response. A 7-gene immune signature (7ISG), related to viral mimicry signalling, is at the core of the HGSC immune response to HKMTi-1-005. In *MYC*-amplified HGSC patients, a low 7ISG score was associated with poor survival. Additionally, *MYC*-amplified cell lines were significantly more sensitive to HKMTi-1-005, whilst a low 7ISG score was associated with greater HKMTi-1-005 sensitivity, effects that were independent of canonical cMYC transcriptional activation. Examining the effects of HKMTi-1-005 treatment in a *MYC*-deregulated lung adenocarcinoma (LuAd) revealed induction of an immune response *in vitro* and prolonged survival *in vivo.* This suggests that inhibition of H3K27/H3K9 methylation maintenance will have efficacy in cMYC-deregulated tumours with low 7ISG scores, via disruption of cMYC-mediated repression of cell autonomous immune signalling and induction of an anti-tumour immune response.

**Statement of significance:** Over 70% of cancers are cMYC-deregulated. We show that inhibition of H3K27/H3K9 methylation maintenance relieves cMYC-dependent immune repression and prolongs survival of animal tumour models, suggesting a novel approach to treating cMYC-deregulated tumours.

## Introduction

cMYC is deregulated in the large majority of cancers, driving hallmark features of cancer including proliferation, apoptotic resistance and angiogenesis (1). Although most often recognised for its role in transcriptional activation, cMYC has also been shown to repress transcription of immune genes in neoplastic cells, preventing expression in a cell-intrinsic manner and therefore promoting immune evasion and resistance to cell death (1–3). Immune targets of cMYC repression include both innate immune signalling, such as the Stimulator of Interferon Genes (STING) pathway, and interferon (IFN) signalling pathways (2,3). The best-characterised repressive mechanism involves a complex formed by cMYC and Myc-interacting zinc finger protein 1 (MIZ1, or Zbtb17) (cMYC-MIZ1 complex), which can both prevent MIZ1 target gene transactivation (3) and instruct repression of target genes (4). However, recent literature has illustrated that a distinct cMYC repression effect is achieved independently of MIZ1, in part via physical interaction between cMYC and the histone methyltransferase enzymes Enhancer of Zeste Homolog 2 (EZH2) (5) and Euchromatic Histone Lysine Methyltransferase 2 (EHMT2 or G9a) (6). Additionally, cMYC exists in a feedback loop with these repressive enzymes, able to enhance expression of both EZH2 and G9a (6,7), whilst EZH2 and G9a have been shown to stabilise and upregulate cMYC respectively (5,8). Hence, the EZH2 and G9a enzymes contribute to cMYC-mediated repression both directly and indirectly.

EZH2 and G9a maintain repressive histone 3 lysine 27 tri-methylation (H3K27me3) (9) and histone 3 lysine 9 mono-, di- and tri-methylation (H3K9me1/2/3) marks respectively (10). Targets of dual H3K27- and H3K9-mediated repression include immune stimulatory genes and the pro-apoptotic mediator IL24 (11,12). Additionally, H3K9 and H3K27 methylation are involved in silencing endogenous retroviruses (ERVs) (13) that, if induced, stimulate a viral mimicry immune response (14). Both epigenetic marks have been implicated in the initiation, progression, metastasis and chemoresistance of multiple tumour types (10,15). Further, EZH2 and G9a are overexpressed in several cancers, with higher expression associated with poorer prognosis (9,10). Whilst cMYC has been described as “undruggable” – due to characteristics such as nuclear localisation and the lack of a binding pocket for small molecule inhibitors (16) – the reversible nature of histone methylation marks has led to interest in targeting them pharmacologically (9,10). As such, targeting the histone methylation marks deposited by EZH2 and G9a may provide a strategy to mitigate aspects of cMYC oncogenicity in addition to combatting widespread pro-tumourigenic epigenetic dysregulation.

Single inhibition of EZH2 or G9a has shown some efficacy in various tumour models (17) and the EZH2 inhibitor (EZH2i) tazemetostat is FDA-approved for treatment of follicular lymphoma and epithelioid sarcoma (17). However, epigenetic inhibitors have yet to demonstrate clinical efficacy in most solid tumours (17). Single inhibition approaches may be limited by physical interaction between the two complexes, and the existence of epigenetic redundancy, such as G9a placing the EZH2-canonical H3K27me3 mark (17). Genetic knock-down and pharmacological inhibition of both methyltransferases has been shown to have greater biological effect than inhibition of either alone and is required for maximal tumour growth inhibition (18), and for induction of apoptosis via IL24 (11). To understand further the interplay between histone methylation and cMYC-driven processes, we have investigated the effects of a dual inhibitor targeting maintenance of both H3K27 and H3K9 methylation marks, HKMTi-1-005 (18).

In this report, we begin with a focus on ovarian high grade serous carcinoma (HGSC), a cancer characterised by frequent cMYC copy number alterations (CNA) and with a clear role for the immune system in prognosis, yet with limited treatment options, poor prognosis, and disappointing response to immunotherapeutic intervention thus far (19,20). We have previously shown that HKMTi-1-005 treatment increased survival of an HGSC model (*Trp53-/-* ID8) (21). This survival benefit was accompanied by immune stimulation including a viral mimicry effect, with expression of ERVs and interferon-stimulated genes (ISGs), alongside increased tumor infiltration by CD8^+^ T effector cells and natural killer (NK) cells and reduced infiltration of FoxP3^+^ T regulatory (Treg) cells (21).

Here, we investigate the hypothesis that immune stimulation in HGSC is achieved in part via disruption of cMYC-mediated repression. This may occur through antagonism of H3K27 and H3K9 methylation, with amplification via induction of viral mimicry. We identify a 7-gene signature (7ISG) that is at the core of the HKMTi-1-005 immune effect and show that this 7ISG score is associated with improved prognosis in patients with *MYC*-amplified HGSC. We illustrate that cell lines with *MYC* amplification are more sensitive to HKMTi-1-005, and that the 7ISG has predictive value for compound sensitivity independent of canonical cMYC activity. Finally, we look to validate HKMTi-1-005 effects in another cancer with frequent *MYC*-deregulation, lung adenocarcinoma (LuAd) (22). In particular, the large subset of LuAd tumours with a *KRAS^G12D^* mutation is evidenced to be dependent on endogenous cMYC activity for tumourigenesis (23), and it has been demonstrated that cMYC suppresses interferon signalling in *KRAS*-driven LuAd models (24). Here we show that treatment with HKMTi-1-005 stimulates immune gene expression in lung cancer lines, and reduces tumour burden whilst prolonging survival in the *KRAS*-driven *MYC*-overexpressing KM mouse model.

## Materials and Methods

### HGSC models: treatment and sequencing

The HGSC cell lines PEO1, PEO4, PEA2, OVCAR5 and OVCAR8 were obtained from Drs Langdon and Melton, Institute of Genetics and Molecular Medicine, Edinburgh University. STR genotyping prior to isolation of RNA authenticated the cell lines. Cells were seeded at a density of 250,000 cells per well in a 6-well plate for 24 hours before treatment with HKMTi-1-005 at their respective IC50 values for 72 hours, ahead of harvesting for sequencing analyses. For RNAseq, RNA extraction with rRNA depletion was performed. Subsequently, paired-end RNAseq was performed by the Institute of Cancer Research (ICR). followed by downstream processing on the raw data generated (see supplementary methods). For ATACseq, harvested cells were processed using the protocol described in (32), before sending samples to the ICR for sequencing. See supplementary methods for downstream processing. Collection and initial processing of ID8 mouse model data are described in (21). ID8 RNAseq and ATACseq data are available via ENA (https://www.ebi.ac.uk/ena/submit/sra/#home), primary accession number PRJEB44851, secondary accession number ERP12894. Processing of raw data is described in supplementary methods.

### Patient cohorts

Initial survival analyses utilised a cohort of 93 HGSC patients, from The International Cancer Genome Consortium (ICGC), accessible at the ICGC Data Portal (https://dcc.icgc.org/releases/current/Projects/OV-AU) (33). FPKM-normalised reads (exp_seq.OV-AU.tsv file) were downloaded from the portal alongside clinical data (donor.OV-AU.tsv file) before filtering and scoring as described in ‘Genesets and Scoring’. A validation cohort consisted of 429 ovarian cancer patients from The Cancer Genome Atlas (TCGA) study. The R package TCGABioLinks (34) was used to download count data as TPM, prior to filtering and scoring for genesets as described below. Samples were annotated with their cMYC CNA status and clinical information using cBioPortal download (28). After filtering to include Grade 2 or 3 patients who had not received radiotherapy or neoadjuvant chemotherapy, 111 samples with a cMYC amplification and 214 without were retained for use in survival analyses. Disease-specific survival was defined by TCGA as the date of initial diagnosis to date of death specifically from disease.

### Sanger pan-tumour cell lines: sensitivity, doubling time, gene expression, mutation and amplification status

Sensitivity data in the form of IC50 values were downloaded from the publicly available resource, The Genomics of Drug Sensitivity in Cancer 2 (GDSC2) (25) (https://www.cancerrxgene.org/downloads/bulk_download, accessed 30/09/22). Methods for calculating IC50 values based on cell viability assays are described in the accompanying documentation available at the same site. Raw data downloaded included sensitivity measurements for 295 compounds, across 969 cell lines derived from many different tumour types. Within these data, there were two single EHMT2/G9ai (UNC0638 and A-366, DRUG IDs 2038 and 2157 respectively), and two single EZH2i (GSK343 in two separate screens, DRUG IDs 1627 and 2037), alongside HKMTi-1-005. Of these, we selected UNC0638 for an EHTM2/G9ai comparison, as an inhibitor with the same substrate-competitive mechanism of action as BIX01294 from which our compound was derived, unlike the peptide-competitive A-366. For EZH2i comparison, we included the GSK343 data with the greatest overlap in cell lines tested as compared to HKMTi-1-005 (DRUG ID 2037).

Cell lines from solid tumours with sensitivity data for all three compounds of interest were included in analyses (n = 565). For these lines, growth rate (doubling time, hours) was obtained from the Cell Model Passports hub hosted by The Sanger Institute (https://cellmodelpassports.sanger.ac.uk/downloads, 07/09/22 release). Where more than one value for a cell line was included in the data, an average of recorded growth rates was taken. Gene expression (in the form of RMA data), mutation status and amplification data were downloaded from the COSMIC Cell Lines Project v96, released 31/05/22. Copy number alteration such as those recorded for cMYC are defined by COSMIC as an amplification if the average genome ploidy <= 2.7 and total copy number >= 5, or the average genome ploidy > 2.7 and the total copy number >= 9.

### Genesets and scoring

Genesets described were downloaded from MSigDB where available (https://www.gsea-msigdb.org/gsea/msigdb/human/genesets.jsp, version 7.4 released 04/21, accessed 20/10/22 and 22/03/23. See Box 1). The equivalent mouse genesets were downloaded from the same site where possible (https://www.gsea-msigdb.org/gsea/msigdb/mouse/genesets.jsp, see Box 1). Where no equivalent geneset existed, AnnotationDbi (https://bioconductor.org/packages/release/bioc/html/AnnotationDbi.html) was used to map human genes to mouse genes. The cMYC-G9a repression geneset was created based on literature published describing a list of 38 genes repression by this complex in breast cancer cell lines (6), whilst the 7-gene ISG signature was created based on ISGs strongly or consistently induced by the HKMTi-1-005 in HGSC cell lines tested (Supplementary figure 5), as described in Results. The dual EZH2/G9a target geneset was defined by Casciello et al. (11). The singscore package was used to score samples for these genesets (https://bioconductor.org/packages/release/bioc/html/singscore.html), based on FPKM (ICGC), TPM (TCGA) or RMA counts (GDSC2) to ensure correction for gene length bias. FPKM/TPM counts were filtered prior to scoring, to ensure > 0.5 FPKM/TPM in more than half the samples. Immune infiltration was estimated from filtered counts, using the CIBERSORTx platform (33) in absolute mode with 100 permutations, batch correction and with quantile normalisation disabled.

#### Box 1.

**Description of genesets used for cell line/sample scoring.**

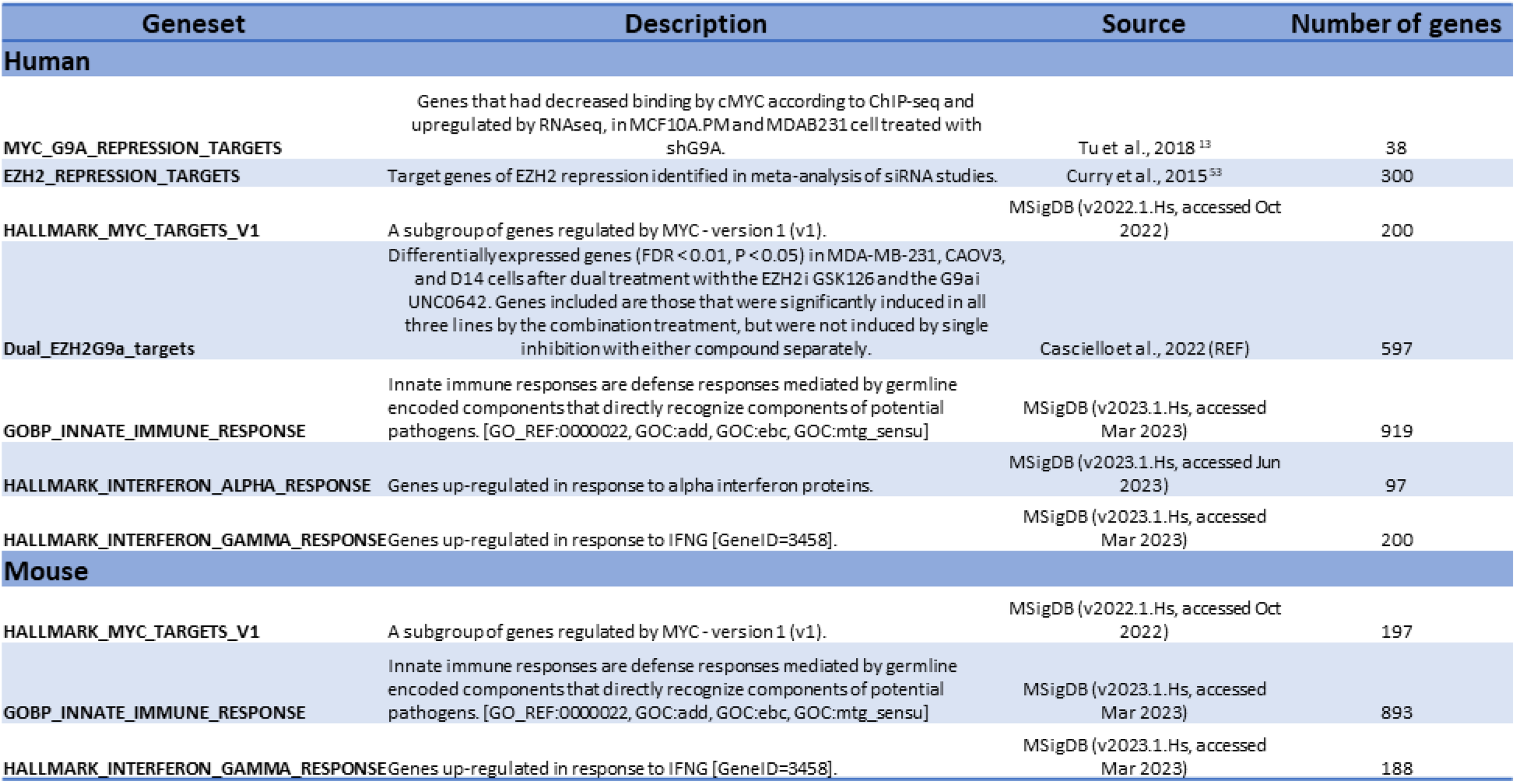

### Statistical analyses

Unless otherwise stated, graphs and statistics were generated in the R environment (4.2.2) (2022-10-31) using key R packages including ggplot2 and tidyverse. If the data were normally distributed, or could be transformed to be so, correlation analyses were performed using a Pearson’s analysis. If non-normal, correlations were performed using a Spearman’s analysis. Groupwise comparisons were performed using t-tests or Mann-Whitney/Wilcoxon tests, for normal or non-normal data respectively, using ggpubr and ggstatsplot packages. Multiple linear regression was performed using the lm function from the R base package stats, with added-variable plots for visualisation generated using the car package.

CoxPH models were generated using the survival package, with testing of the Cox proportional hazards assumptions and variance inflation factors using the survival and car packages. The age variable was stratified at 50 years old as used elsewhere (34). Kaplain-Meier curves were used to visualise the effect of scores on survival, splitting into high and low groups at the median 7ISG value for the cohort plotted. Statistical analyses in lung cell lines and mouse models were performed using GraphPad Prism.

### KM LuAd mouse model and in vivo procedures

Procedures involving mice were performed in accordance with Home Office Licence number PE47BC0BF (CRUK BICR, UK). The *LSL-KRas^G12D^* (RRID:IMSR_JAX:008179) and Rosa26^DM.lsl-MYC^ (RRID:IMSR_JAX:033805) allelic mice were described previously (30,46). Mice were maintained on a mixed FVBN/C57Bl6 background, housed on a 12-hour light/dark cycle, and fed and watered ad libitum. Recombinant Adenovirus expressing CRE recombinase, Ad5mSPC-CRE, was purchased from the University of Iowa gene therapy facility. For Ad5mSPC-CRE instillation, young adult mice (8- to 10-week-old) were sedated with a mixture of medetomidine and ketamine, injected IP. For all experiments, 1×10^8^ pfu Ad5mSPC-CRE were administered intranasally using the calcium phosphate precipitation method, as described previously (47). Mice were randomly assigned to treatment groups, balanced only for sex, and both males and females were used in all studies. HKMTi-1-005 (40mg/kg) or vehicle control (0.9% NaCl) was administered twice daily by IP injection, from 4-weeks post induction to 8-weeks post induction, with a treatment schedule of three days treatment followed by 4 days recovery each week. To assess the potential survival benefit of treatment, HKMTi-1-005 (35mg/kg) or vehicle control (0.9% NaCl) was administered twice daily by IP injection for 3 consecutive days/week for 4 weeks, as above, and mice were monitored by facility staff without knowledge of experimental parameters. Mice were humanely culled when clinical endpoint was reached.

### KM LuAd mouse model: immunohistochemistry and tissue analysis

All mouse tissue IHC staining was performed on 4µm, formalin-fixed, paraffin-embedded sections, which had been previously heated to 60°C for 2 hours. Peroxidase blocking as performed for 10 minutes in 1% H_2_O_2_ diluted in H_2_O, followed by enzyme-mediated antigen retrieval. Nonspecific antibody binding was blocked with 1% BSA for one hour at room temperature. Ki67 (Cell Signalling 12202) was used at a dilution of 1:1000, with Rabbit Envision (Aligent) used as secondary antibody. The horseradish peroxidase (HRP) signal was detected using liquid DAB (Aligent and Invitrogen). Sections were counterstained with hematoxylin and cover-clipped using DPX mount (CellPath). Tumour burden and Ki67+ cells were quantified using HALO Software (Indica Labs).

### Lung cancer cell lines and treatment

A549 & H358 cells were obtained from ATCC and were maintained in RPMI supplemented with 10% FBS and penicillin-streptomycin. Cell lines were thawed from primary stocks maintained under liquid nitrogen and cultured for a maximum of 8 weeks (<20 passages), during which time all experiments were performed. Dose response curves for HKMTi-1-005 treatment were generated by Incucyte analysis. For analysis of IFN regulator expression, cells were treated with 10µM HKMTi-1-005, 100nM Trametinib, or H_2_O (control) for 24 hours and harvested for RNA analysis. Total RNA was extracted from samples using Qiagen RNeasy kit as per manufacturer’s protocol. DNA was depleted using the RNase-Free DNase Set (Qiagen). In the final step, total RNA was eluted in 30µl of RNase-Free H_2_O. The concentration of RNA was determined using NanoDrop 2000c (Thermo Fisher). cDNA was generated using the QuantiTect Reverse Transcription kit (Qiagen), using 2µg RNA in each reaction. Expression of genes of interest was quantified by real-time PCR using SYBR Green method (VWR QUNT95072). Primers used are detailed in supplementary methods. Expression of Interferon-related genes was normalised to expression of GusB.

### Data availability

ID8 RNAseq and ATACseq data are available via ENA (https://www.ebi.ac.uk/ena/submit/sra/#home), primary accession number PRJEB44851, secondary accession number ERP12894. PEO1, PEO4, PEA2, OVCAR5 and OVCAR8 RNAseq and ATACseq data are available from the corresponding author and are in process of being made publicly available.

### Ethics statement

Procedures involving mice were performed in accordance with Home Office Licence number PE47BC0BF (CRUK BICR, UK) and approved by Glasgow University Animal Ethics committee. Access to anonymised patient data were approved by the respective study access committees and study specific ethics.

## Results

### Greater cMYC-G9a- and EZH2-mediated repression correlates with lower immune scores, an effect heightened in cMYC-amplified cell lines

The Genomics of Drug Sensitivity in Cancer 2 (GDSC2) is a publicly available dataset, describing drug sensitivity, gene expression and copy number alteration (CNA) amongst other cellular characteristics in a pan-tumour collection of cell lines (25). Geneset scoring was used to assess the transcriptional profiles of 565 cell lines derived from solid tumours (see Methods). Cells were scored for genesets reflecting the extent of hallmark cMYC transcriptional activity, cMYC-G9a- and EZH2-mediated repression, and immune activity (Box 1). These data revealed a strong positive correlation between lower cMYC-G9a or EZH2 repression and higher immune signalling, including both innate immune and interferon alpha or gamma signalling (Fig 1A, r > 0.48, p < 0.001). The opposite was observed when comparing hallmark cMYC activity scores with immune scores, where greater transcriptional activity at cMYC hallmark genes strongly anticorrelated with immune signalling (Fig 1A, r = −0.45, p < 0.001).

**Figure 1.**
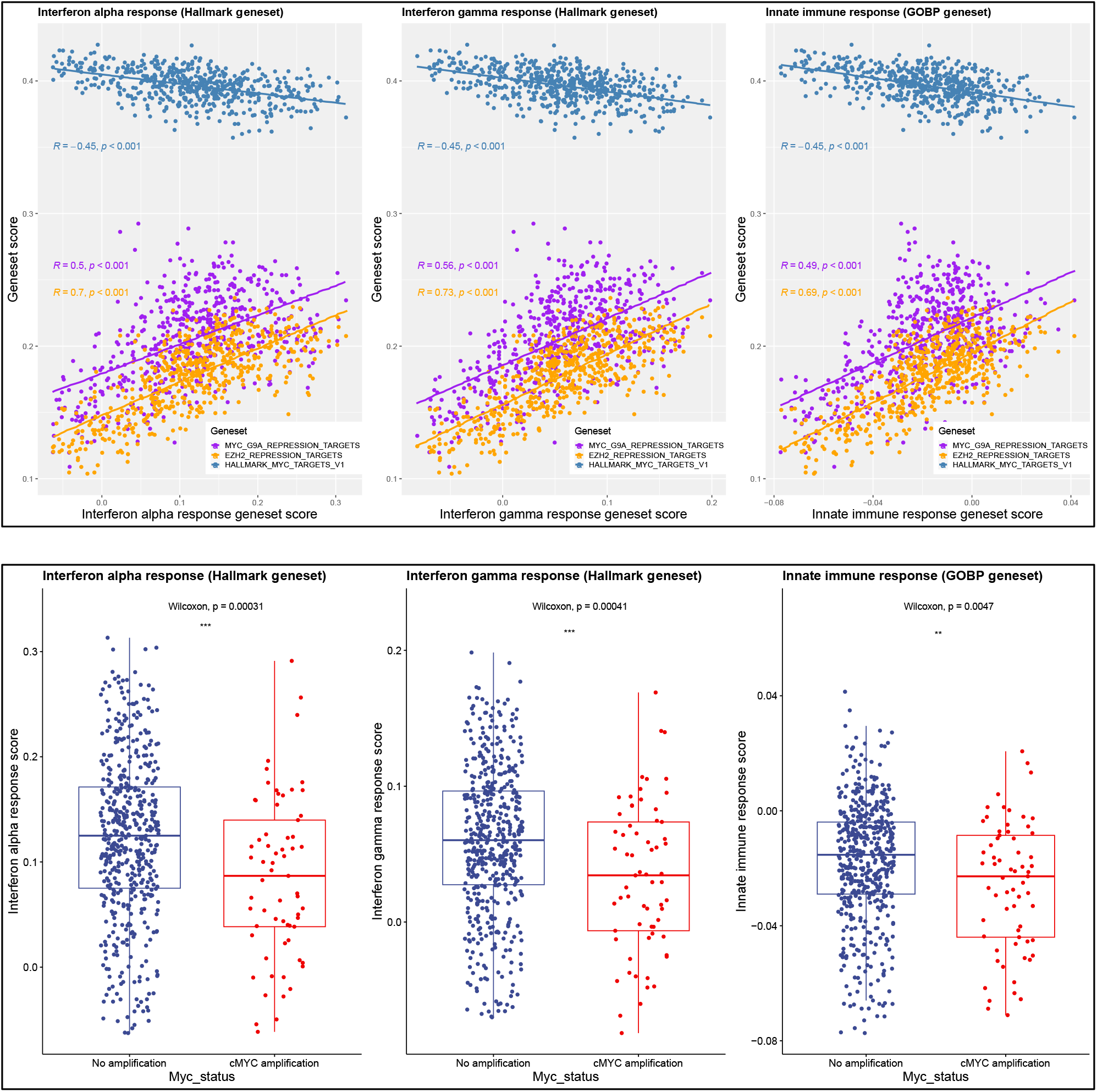
Greater cMYC-G9a/EZH2-mediated repression correlates with lower immune scores. (A) Spearman’s correlation between immune genesets (x axis) and genesets reflecting targets of cMYC-G9a repression, EZH2 repression, and hallmark cMYC activation (y axis), in solid tumour cell line panel (n = 565). (B) Comparison of immune geneset scores between cMYC-amplified (n=65) and non-amplified (n=500) cell lines (Wilcoxon t-test).

We next examined whether *MYC*-amplified cell lines demonstrated a stronger cMYC-mediated repression effect in this panel. As expected, *MYC*-amplified lines (n=65) demonstrated greater canonical cMYC activity, illustrated by higher scores for the hallmark cMYC activity geneset (Supplementary figure 1, Box 1). However, these lines also showed greater repression by cMYC-G9a/EZH2 (Supplementary figure 1), and lower levels of immune signalling than those without an amplification (n=500) (Fig 1B). These data support the hypothesis that cMYC represses innate immune and interferon targets in a cell-intrinsic manner, including via collaboration with histone methylation marks deposited by G9a and EZH2.

### HKMTi-1-005 is an epigenetic inhibitor with dual activity against H3K27 and H3K9 methylation

Genetic inactivation of EZH2 and G9a together results in greater inhibition of tumour cell growth and synergistic induction of apoptosis compared with inactivation of either gene alone, alongside transcriptional induction of additional genes and genomic elements (11,18). Based on the quinazoline template of the G9a inhibitor BIX-01294, we have previously identified compounds that reduce methylation of the EZH2/G9a marks H3K27 and H3K9, and induce transcriptional responses consistent with this dual inhibition (18,26). We have focussed on HKMTi-1-005 (Supplementary figure 1B) based on its efficacy for reducing H3K27/H3K9 methylation levels and inhibition of cell growth *in vitro* and specificity for EZH2 and G9a inhibition over other methyltransferases (18).

To examine the effects of HKMTi-1-005 on H3K27- and H3K9-mediated transcriptional regulation further, five HGSC cell lines were investigated. PEA2, OVCAR5, OVCAR8, PEO1 and PEO4 were treated with HKMTi-1-005 at their respective IC50 values and gene expression assessed 72h later by RNAseq. Supporting the existing literature describing IL24 as a target of dual repression by H3K27 and H3K9 methylation based on genetic inactivation of EZH2 and EHMT2/G9A (11), we observed significant upregulation of *IL24* in 4/5 cell lines (p-adjusted < 0.05, Fig 2A). Geneset enrichment analysis (GSEA) further demonstrated a significant change in the EZH2/G9a target geneset curated by Casciello et al., in all cell lines (Box 1, Fig 2B). This geneset comprises genes that are differentially expressed (both up- and down-regulated) only following simultaneous inhibition of both EZH2 and G9a, but not with inhibition of either individually (11) (Box 1, Fig 2B). To examine the effect of HKMTi-1-005 on EZH2/G9a targets *in vivo*, previously generated RNAseq data derived from *Trp53−/−* ID8 ovarian tumours (21) were re-analysed. These data, comparing untreated and HKMTi-1-005-treated mice (21), once again showed upregulation of *IL24* with 4.3 fold-change (p-value 0.02, Fig 2A) and significant enrichment of the EZH2/G9a geneset following HKMTi-1-005 treatment (Fig 2B).

**Figure 2:**
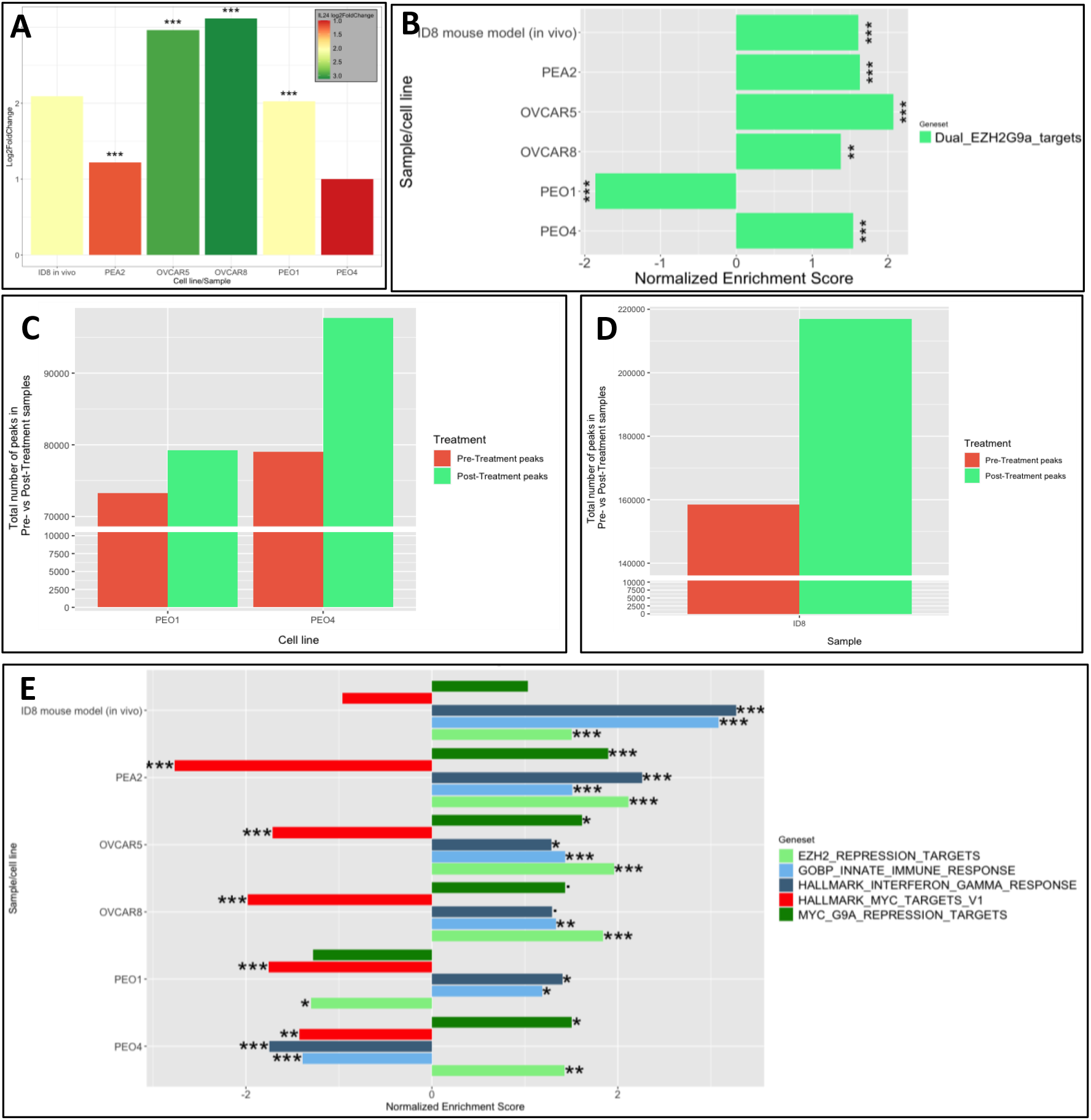
HKMTi-1-005 is an inhibitor of H3K27 and H3K9 that antagonises cMYC effects. (A) Log2 fold changes and associated p-adj values of IL24 gene expression after HKMTi-1-005 treatment in HGSC cell lines PEA2, OVCAR5, OVCAR8, PEO1 and PEO4, alongside tumour samples from the ID8 *in vivo* murine HGSC model. (B) Geneset enrichment analysis showing changes in a dual EZH2/G9a target geneset. (C) Number of peaks of chromatin accessibility in pre- vs post-HKMTi-1-005 treatment HGSC cell lines PEO1 and PEO4, based on ATACseq analysis. Scale break added at 10,000 to 70,000 peaks. (D) Number of peaks of chromatin accessibility in pre- vs post-HKMTi-1-005 treatment ID8 tumour-derived samples, based on ATACseq analysis. Scale break added at 10,000 to 140,000 peaks. (E) Geneset enrichment analysis plot of changes in targets of cMYC-G9a and EZH2 repression, innate immune, interferon gamma and hallmark cMYC genesets after HKMTi-1-005 treatment. . = p-adj < 0.1, * = p-adj < 0.05, ** = p-adj < 0.01, *** = p-adj < 0.001.

An assay for transposase-accessible chromatin with sequencing (ATACseq) was performed on the PEO1 and PEO4 cell lines and the ID8 tumours following HKMTi-1-005 treatment, to confirm the effect on chromatin accessibility. These data confirmed that there were a greater number of accessible chromatin peaks after treatment in each sample (Fig 2C, Fig 2D), consistent with a reduction in the repressive H3K27 and H3K9 methylation marks. Taken together, the findings demonstrate that HKMTi-1-005 has EZH2 and G9a inhibitory-like effects on gene expression and chromatin accessibility.

### HKMTi-1-005 relieves repression by cMYC-G9a/EZH2 whilst downregulating canonical cMYC activity and inducing an immune response in HGSC

We next examined genesets reflecting cMYC-G9a or EZH2 repression, hallmark cMYC transcriptional activity and immune responses in HKMTi-1-005-treated HGSC cell lines and ID8 tumours. Whilst upregulating cMYC-G9a/EZH2-repressed genes, HKMTi-1-005 stimulated a downregulation in hallmark cMYC transcriptional activity (Box 1, Fig 2E). The upregulated genesets varied by sample, suggesting some variability in targets of cMYC-G9a/EZH2 repression depending on cell line characteristics, as expected (6,18). However, all lines showed significant induction of genesets consistent with relief of cMYC-G9a/EZH2 repression (Supplementary figure 2A-D).

Innate immune and interferon gamma response genesets were consistently upregulated after treatment, with the strongest effect *in vivo* (Fig 2E). This was not true of interferon alpha signalling, which showed variable responses between cell lines (Supplementary figure 3A). Of note, PEO4 cells – derived from the same patient as PEO1, but after resistance to cisplatin had emerged – did not replicate this trend, with downregulation of both immune genesets. PEO4 demonstrated immune signalling genesets prior to treatment that were 2 to 4-fold higher than other lines (Supplementary figure 3B), suggesting a possible cause for this lack of induction.

### Immune stimulation by HKMTi-1-005 includes a viral mimicry response, and is characterised by a 7-gene signature (7ISG)

Given literature describing a role for EZH2 and G9a in silencing transposable elements (TEs) that can stimulate a viral mimicry immune response (13,14), we investigated TE changes after HKMTi-1-005 treatment. Subfamily- and locus-level analyses showed TEs were upregulated in all samples after treatment (Supplementary figure 4A-D). This included a consistent upregulation of ERVs, the best-described inducers of viral mimicry (Fig 3A, Supplementary figure 4E). Analyses of ID8 tumours strengthened evidence for HKMTi-1-005-induced upregulation of ERVs (Fig 3C, Supplementary figure 4F), with a greater absolute number of induced ERVs than in cell lines.

**Figure 3:**
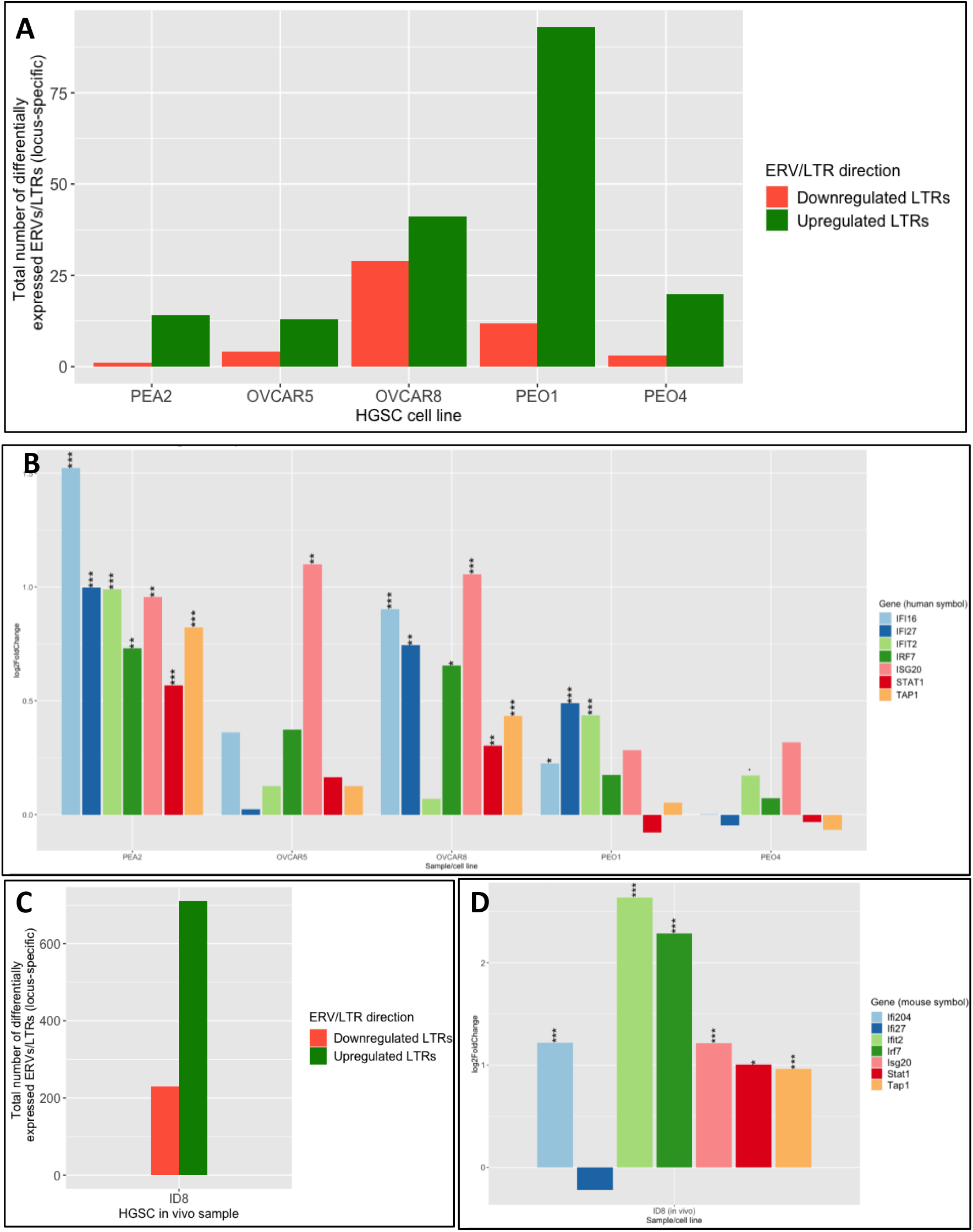
HKMTi-1-005 induces a viral mimicry effect in vitro and in vivo, characterized by the 7ISG. (A) Differentially expressed ERVs/LTRs across HGSC cell lines after treatment with HKMTi-1-005, at a locus-specific level. (B) Changes in expression of 7 ISGs across HGSC cell lines after HKMTi-1-005 treatment. (C) Differentially expressed ERVs/LTRs in ID8 tumours after treatment with HKMTi-1-005, at a locus-specific level^+^. (D) Changes in expression of 7 ISGs in ID8 tumours after HKMTi-1-005 treatment^+^. ^+^ ID8 data are an updated analysis of data published in Spiliopoulou et al., 2022, with re-mapping of raw data to an updated genome and a novel methodology for TE analysis. . = p-adj < 0.1, * = p-adj < 0.05, ** = p-adj < 0.01, *** = p-adj < 0.001.

Previously, Chiappinelli et al. illustrated induction of viral mimicry using a DNA demethylating agent (14), defining 28 interferon-stimulated genes (ISGs) when examining the viral mimicry response in ovarian tumours (27). We examined differential expression of these genes following HKMTi-1-005 treatment and identified a core 7-gene signature (7ISG) reflecting genes that were strongly or consistently upregulated by the compound (Fig. 3B, Supplementary figure 5A). Induction of the 7ISG was confirmed in ID8 tumours, with 6/7 genes significantly upregulated after treatment (p-adjusted <= 0.05) (Fig 3D). Taken together, these data suggest that HKMTi-1-005 is capable of relieving cMYC-G9a/EZH2-mediated repression in HGSC cell lines and *in vivo* models, whilst inducing an immune response, including viral mimicry, that is defined by the 7ISG gene signature.

### *In vitro* sensitivity to HKMTi-1-005 is greater in *MYC-*amplified lines and significantly associated with the 7ISG, independent of canonical cMYC activity

Having established that HKMTi-1-005 relieves cMYC-mediated repression whilst inducing an immune response, we hypothesised that cMYC status and immune/7ISG scores at baseline might relate to cell autonomous compound sensitivity. Sensitivity to HKMTi-1-005 in the GDSC2 solid tumour cell line panel (n=565) was compared to an EZH2 inhibitor (GSK343) and a G9a inhibitor (UNC0638).

Lines with *MYC* amplification – which showed greater cMYC-G9a/EZH2 repression (Fig 1B) – were significantly more sensitive to the HKMTi-1-005, independent of doubling time (Fig 4A, Supplementary figure 6A, 6B). These *MYC*-amplified lines were also more sensitive to G9ai and EZH2i than non-amplified lines (Fig 4A). In *MYC*-amplified lines, there was a strong correlation between lower immune/7ISG scores and greater sensitivity to both HKMTi-1-005 and the single G9ai, but not to EZH2i (Fig 4B). Conversely, higher canonical cMYC signalling correlated with greater sensitivity to all compounds. Non-amplified lines demonstrated these correlations between sensitivity and immune/7ISG/hallmark cMYC scores to a lesser extent, but with similar trends (Fig 4C). Given that 7ISG and hallmark cMYC scores significantly anti-correlate (Supplementary figure 6C), and that both scores correlate with sensitivity to the compounds, we looked to investigate the individual contribution of each score to compound sensitivity.

**Figure 4:**
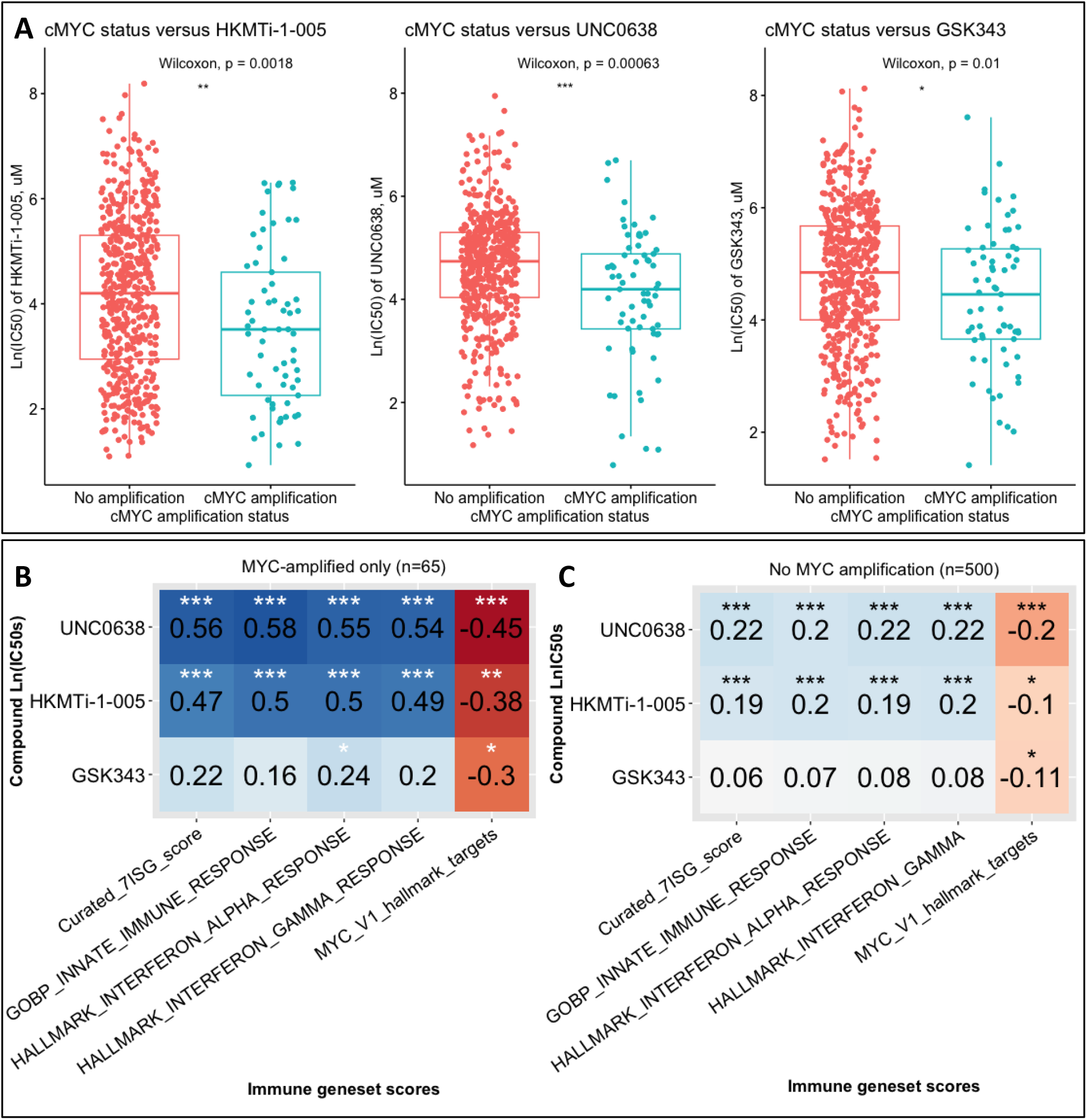
Sensitivity to HKMTi-1-005 relates to cMYC status and 7ISG scores. (A) Difference in sensitivity to HKMTi-1-005 or cisplatin between cMYC-amplified (n=65) and non-amplified (n=500) solid cell lines (Wilcoxon test). (B) Spearman’s correlation between sensitivity to compounds and geneset scores across cMYC-amplified (n=65) and non-amplified (n=500) solid cell lines. Compounds include (top to bottom) UNC0638 (single G9ai), HKMTi-1-005, GSK343 (single EZH2i) and cisplatin. Genesets include (left to right) the 7ISG, innate immune response, interferon alpha response, interferon gamma response and hallmark cMYC targets. * = p < 0.05, ** = p < 0.01, *** = p < 0.001

To do so, we used a multiple linear regression approach. In each case, drug sensitivity was the dependent variable, whilst 7ISG and hallmark cMYC scores were independent variables. This analysis indicated that sensitivity to HKMTi-1-005 in both *MYC*-amplified and non-amplified lines was significantly associated with the 7ISG score, independent of hallmark cMYC activity (Supplementary figure 7A, 8A). Hallmark cMYC activity was not independently associated with HKMTi-1-005 sensitivity in either cohort. By contrast, both hallmark cMYC activity and the 7ISG were independent predictors of G9ai sensitivity in *MYC*-amplified and non-amplified lines (Supplementary figure 7B, 8B). There was no significant association between EZH2i sensitivity and the 7ISG score in either cohort (Supplementary figure 7C, 8C). These data support the hypothesis that the 7ISG, but not hallmark cMYC activity, is an independent predictor of HKMTi-1-005 sensitivity in *MYC*-amplified and non-amplified lines, and that this relationship is a distinct feature of HKMTi-1-005 sensitivity as compared to single G9ai or EZH2i.

### The HKMTi-1-005-induced 7ISG signature is associated with improved prognosis in *MYC*-amplified HGSC

To investigate the clinical relevance of the 7ISG, we examined a cohort of HGSC patients obtained from the International Cancer Genome Consortium (ICGC, n = 93). RNA data were scored for the 7ISG (see Methods). Immune infiltration for each sample, as inferred using CIBERSORTx, indicated that the 7ISG positively correlated with several immune cell types (Fig 5A). This included CD8^+^ T cells, which are associated with improved prognosis in HGSC (19), as well as activated natural killer cells, dendritic cells and gamma delta T cells. Moreover, 7ISG was associated with a significantly longer overall survival in both uni- and multi-variable analyses (p = 0.05 and p = 0.03 respectively, Fig 5B).

**Figure 5:**
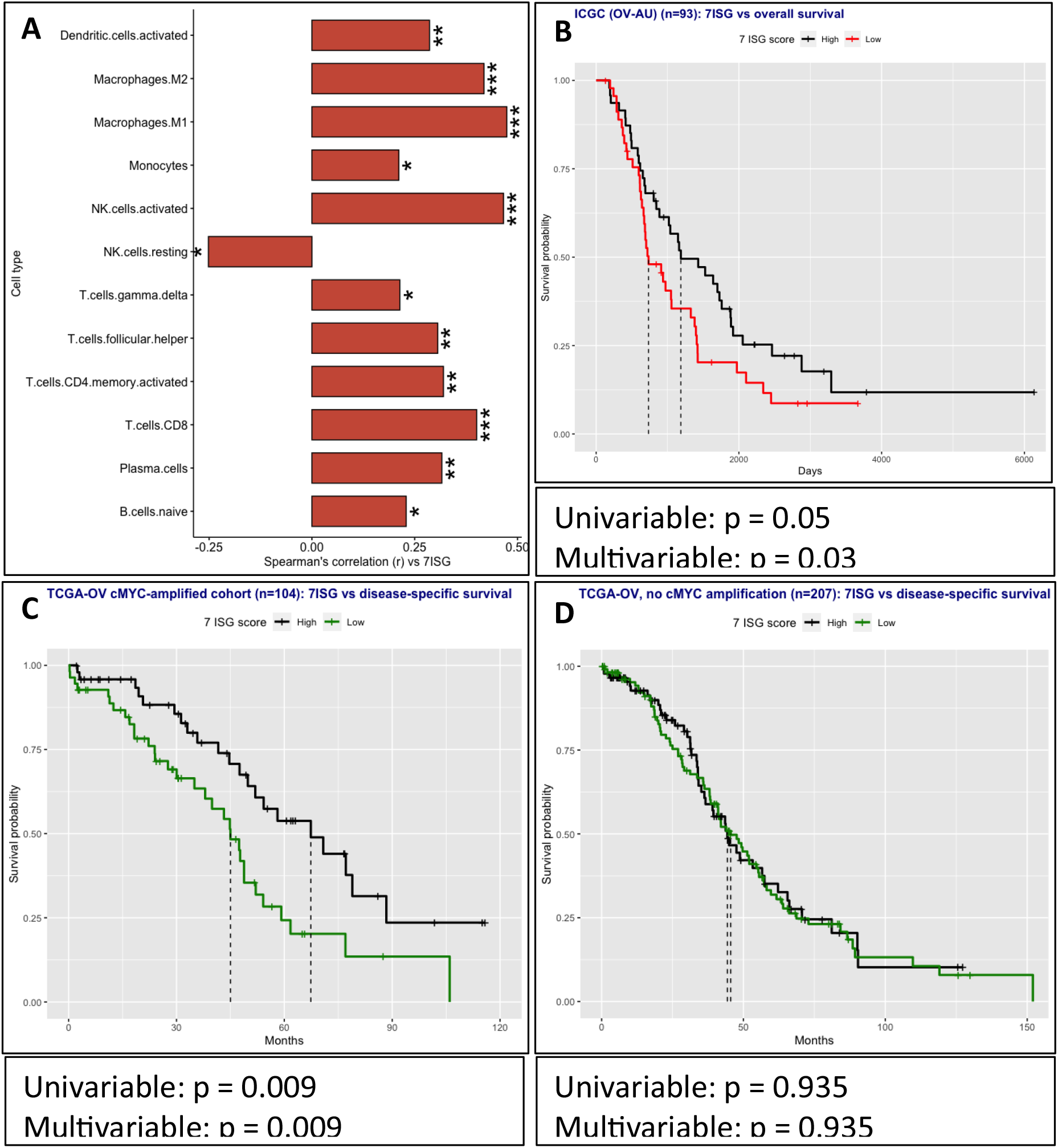
A higher 7ISG is associated with improved prognosis in cMYC-amplified HGSC patients. (A) Spearman’s correlation between 7ISG scores and estimates of immune infiltration, in a cohort of HGSC patients (n=93) – significant associations only, p-value < 0.05. (B) Kaplain-Meier (KM) survival curve and Cox proportional hazards results illustrating difference in overall survival between high and low 7ISG score HGSC patients (n=93). (C) KM survival curve and Cox proportional hazards results illustrating difference in disease-specific survival between high and low 7ISG score patients, in an independent cohort of HGSC patients with a cMYC amplification recorded (n=104). (D) KM survival curve and Cox proportional hazards results illustrating difference in disease-specific survival between high and low 7ISG score patients, in an independent cohort of HGSC patients without a cMYC amplification recorded (n=207). * = p < 0.05, ** = p < 0.01, *** = p < 0.001. For Kaplain-Meier survival curves, scores are stratified into high and low at the median 7ISG value for each cohort. Vertical dashed lines represent median survival for high and low score groups.

Given the effect of *MYC* amplification on cMYC-G9a/EZH2-mediated repression and immune scores, we assessed whether *MYC* amplification influences the prognostic capacity of the 7ISG using a larger independent cohort of HGSC patients from the TCGA database (TCGA-OV) (n=429). RNA data for these samples were scored for the 7ISG and annotated with their *MYC* copy number status and clinical information using cBioPortal (28).

In *MYC*-amplified samples, there was a significant relationship between a higher 7ISG and longer disease-specific survival (univariable p = 0.009, multivariable p = 0.009) with a clear separation between high- and low-score patients (Fig 5C). Interestingly, in HGSC without a *MYC* amplification, the 7ISG had no prognostic value for disease-specific survival (p = 0.9) (Fig 5D). Hence, the 7ISG has prognostic value in HGSC patients with a cMYC amplification, but not those without.

### Treatment of a *MYC*-deregulated lung adenocarcinoma model with HKMTi-1-005 reduces tumour burden and prolongs survival

Having previously demonstrated the sensitivity of *MYC*-amplified cell lines to HKMTi-1-005 and evidenced its therapeutic benefit in a murine model of HGSC, we looked to further validate HKMTi-1-005 efficacy in another *MYC*-deregulated cancer type, lung adenocarcinoma (LuAd) (22). *KRAS*^G12D^-driven LuAd has been shown to be cMYC-dependent (23), and cMYC has been shown to mediate suppression of interferon genes in this subset of LuAd (24). Further, in a LuAd model with *KRAS*^G12D^ and co-expression of APOBEC3B and cMYC, it has been demonstrated that HKMTi-1-005 treatment induces an immune response including upregulation of at least 5 of the 7 genes in the 7ISG score, whilst downregulating proliferation and cell cycle genes (29). HKMTi-1-005 treatment of human lung cancer cell lines *in vitro* suppressed proliferation and induced expression of key interferon regulators, *IRF7*, *IRF9* and *STAT2* (Fig. 6A, B).

**Figure 6.**
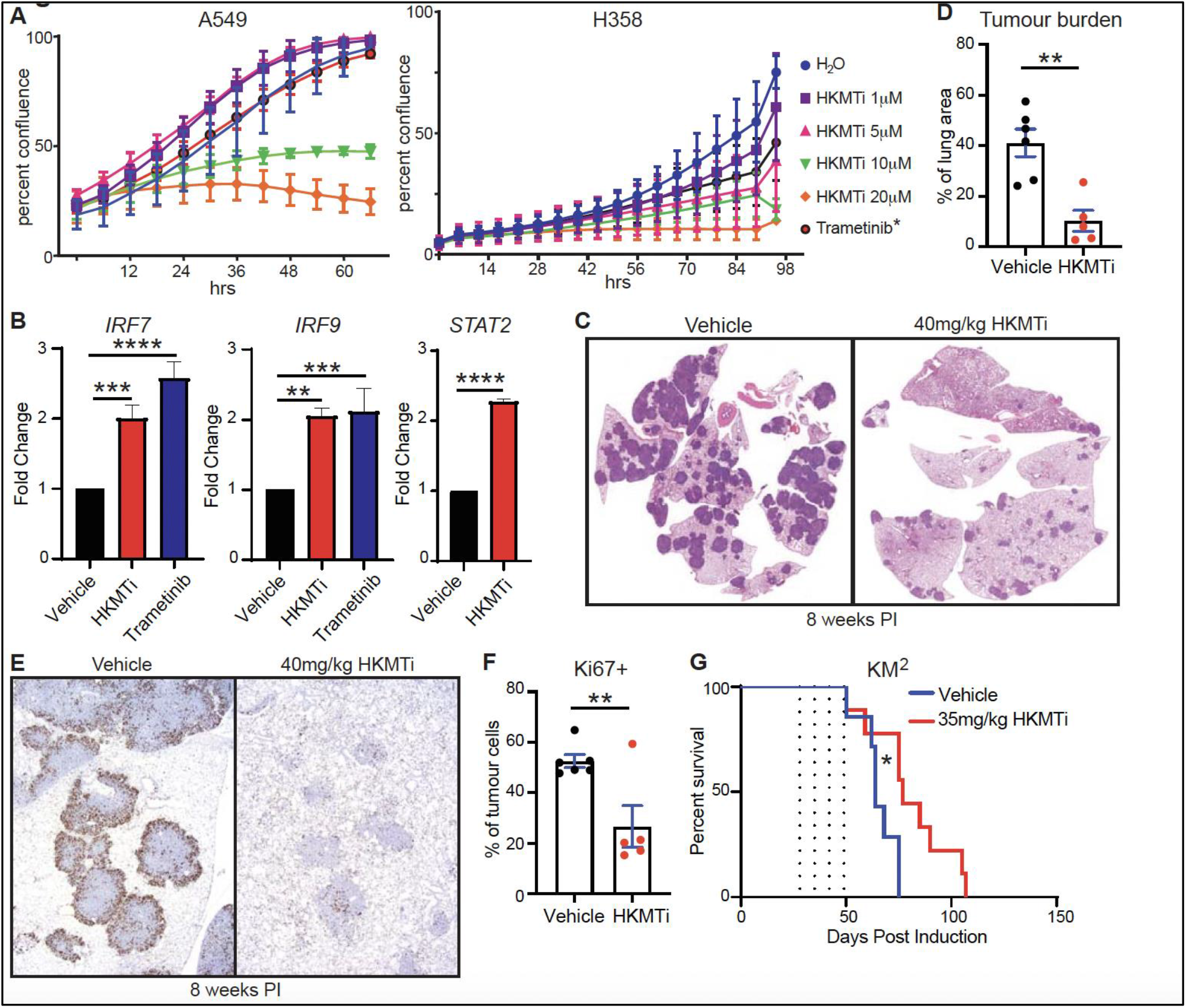
HKMTi-1-005 reduces tumour burden and increases survival in the KM^2^ model of LuAd. **A)** Incucyte analysis of A549 and H358 cell propagation following treatment with the indicated doses of HKMTi-1-005 or H_2_O vehicle. Trametinib positive control was used at 100nM (A549) or 10nM (H358). Error bars reflect SD of 3 biological replecates performed on separate days. P<0.001 for A549 treated with 10 or 20mM HKMTi-1-005; p<0.001 for H358 treated with 5, 10 or 20mM HKMTi-1-005 (2-way ANOVA). **B)** Q-PCR of IFN-related gene expression in A549 cell lines treated ± HKMTi-1-005 or Trametinib. Mean and SD shown for N = 3 technical replicates, representative of 2 biological repeats. ** denotes *P* <0.01, *** denotes *P* < 0.001, **** denotes *P* <0.0001. **C)** Representative images show tumour burden histology, H&E, of KM^2^ mice treated with Vehicle or HKMTi from 4- to 8-weeks post induction and culled following the last of 4 treatment cycles. **D)** Quantification of overall tumour burden calculated as a percentage of total lung area in KM^2^ mice treated with vehicle (N=6) or HKMTI-1-005 (N=5) from 4- to 8-weeks post induction and culled following the last treatment cycle. Mean ± SEM shown (unpaired t-test). **E)** Representative images of Ki67+ stained tissue sections from KM^2^ mice treated with Vehicle (N=6) or HKMTi-1-005 (N=5) from 4- to 8-weeks post induction and culled following final treatment. **F)** HALO automated quantification of percentage of cells in tumours that are Ki67+. Each data point represents the average score of all tumours in a section from an individual mouse. Mean ± SEM shown (unpaired t-test). ** denotes *P* <0.01. **G)** Overall survival of KM^2^ mice treated with vehicle (N=7) or HKMTi-1-005 (N=9). Dotted lines indicate the first day of each cycle of three days treatment. Mantel-Cox log rank test. * denotes *P* <0.05.

The KM model of sporadic LuAd combines conditional overexpression of cMYC from the *Rosa26* locus with CRE-dependent expression of endogenously expressed KRas^G12D^, and exhibits a *MYC* dosage-dependent rate of progression to adenocarcinoma (30,31). Lung tumours were initiated in adult KM^2^ mice (genotype: *lsl-KRas^G12D^;Rosa26^DM.lsl-MYC/lsl-MYC^*) by intranasal installation of adenovirally expressed CRE recombinase. Mice were randomly assigned to treatment groups and treated with HKMTi-1-005, or vehicle, for 3 days/week x 4 weeks, commencing at 4 weeks post tumour initiation. Lungs harvested immediately following the last treatment showed a dramatic suppression of lung tumour burden and a significant reduction in tumour cell proliferation following treatment with HKMTi-1-005 (Fig. 6C-F). A separate cohort of mice treated with the same dosing schedule and maintained following treatment cessation showed significantly enhanced survival following treatment with HKMTi-1-005 (Fig. 6G). We conclude that HKMTi-1-005 demonstrates efficacy against *MYC*-overexpressing lung tumours.

## Discussion

The histone methyltransferases EZH2 and G9a contribute to carcinogenesis and chemoresistance in many tumour types, by repressing gene expression through methylation of histones at H3K27 and H3K9 (9,10). EZH2 and EHMT2 are frequently amplified or overexpressed in a variety of tumours (18), and higher expression is associated with poor prognosis (9,10). Alongside contributions to global epigenetic dysregulation, both EZH2 and G9a are known to interact with the onco-protein cMYC, which is aberrantly regulated in the majority of tumours (5,6). In addition to transcriptional activation, cMYC drives cell-autonomous repression of immune gene targets (1- 3), in part via recruitment of EZH2/G9a-deposited repressive H3K27 and H3K9 methylation marks (5,6). This interaction is further enhanced by cMYC and EZH2/G9a sitting in a positive feedback loop (5–8). Despite deregulation of cMYC in more than 70% cancers (1), it remains difficult to target therapeutically (16). Disruption of H3K27/H3K9 methylation may provide a novel therapeutic approach, allowing antagonism of cMYC-mediated immune repression, alongside targeting of global EZH2/G9a-mediated epigenetic dysregulation.

EZH2 and G9a inhibitors targeting either enzyme individually have had limited clinical success, especially in solid tumours (17), an observation that may be explained by epigenetic redundance and crosstalk between the two complexes (17). A dual EZH2/G9a targeting approach has been shown to be more effective than single enzyme inhibition for induction of tumour cell death and antitumour immune responses in experimental models (11,18,21). Yet, the clinical and pharmacodynamic evaluation of two single EZH2 and G9a inhibitors is more challenging than evaluation of a single compound with dual activity. HKMTi-1-005 has dual activity against the maintenance of H3K27 and H3K9 methylation marks, and extends survival of HGSC mouse models with accompanying immune stimulation and cell death (21). Here, we have used HKMTi-1-005 to evaluate disruption of cMYC-associated transcriptional networks and induction of an immune response via inhibition of H3K27me/H3K9me maintenance.

We demonstrate that the extent of repression by cMYC-G9a/EZH2 anticorrelates with the extent of immune signalling across 565 pan-cancer solid tumour cell lines, and particularly in cell lines with a *MYC* amplification. In HGSC cell lines and mouse models, HKMTi-1-005 induces changes in gene expression and chromatin accessibility that support the dual inhibition of H3K27me/H3K9me maintenance. This includes induction of *IL24*, described as a target of dual EZH2/G9a repression and capable of inducing apoptosis in a cell-autonomous manner (11). Further, HKMTi-1-005 relieves cMYC-G9a/EZH2 repression whilst downregulating canonical cMYC activity at genes associated with cell proliferation. Accompanying cMYC-G9a/EZH2 de-repression, HKMTi-1-005 induces an immune response in HGSC, with viral mimicry, as characterised by induction of endogenous retroviruses (ERVs) and interferon-stimulated genes (ISGs). A 7-gene signature – 7ISG – is at the core of this response.

Sensitivity to HKMTi-1-005 is higher in *MYC*-amplified lines and significantly associated with lower basal 7ISG scores. This relationship is independent of canonical cMYC transcriptional activation. In addition, higher 7ISGs are associated with improved prognosis in *MYC*-amplified HGSC patient samples. Extending analyses to another *MYC*-deregulated tumour type – LuAd – demonstrates that treatment of lung cancer cell lines with HKMTi-1-005 induces immune gene expression whilst reducing proliferation, suggesting a similar HKMTi-1-005 effect on cMYC-mediated transcriptional repression and activation as observed in HGSC models. Further, we show that in the *in vivo* KM LuAd model, with cMYC overexpression accompanying KRAS^G12D^ expression, HKMTi-1-005 reduces tumour burden and improves survival. These data suggest HKMTi-1-005 may be particularly effective in *MYC*-deregulated tumours with low 7ISG scores, via disruption of the cMYC-mediated immune repression driven by H3K27/H3K9 methylation.

The current correlative study of cMYC-mediated repression of immune signalling and involvement of EZH2/G9a across pan-tumour cell lines supports biochemical observation in experimental models (1-3,5-8). However, further clarity on the exact mechanism of action of HKMTi-1-005 on EZH2 and the cMYC-EZH2 axis is required. For example, the lack of a published cMYC-EZH2 geneset prevents distinction between HKMTi-1-005 effects on single EZH2 target genes versus dual cMYC-EZH2 repression targets, in contrast to the published cMYC-G9a geneset (6) which allows this distinction for G9a effects. Additionally, although HKMTi-1-005 is based on modification of a G9a inhibitor to have EZH2i-like effects, and clearly has both G9ai- and EZH2i-like effects at the transcriptional level in HGSC models, the potency of HKMTi-1-005 on EZH2 enzymatic activity in vitro is weak (18). Potentially, the EZH2i-like effects of HKMTi-1-005 in HGSC models may stem from disruption of the crosstalk between G9a and EZH2 as opposed to direct enzymatic inhibition.

HKMTi-1-005 has been shown to extend survival of *in vivo* HGSC (21) and LuAd models. Although the effects on survival are moderate in these models, the compound is well-tolerated and prolonged treatment may extend this benefit, alongside combination approaches with conventional chemotherapies or immunomodulators. More *in vivo* studies will also be vital to confirm whether 7ISG scores can be used to evaluate HKMTi-1-005 sensitivity and the potential of 7ISG score as a pharmacodynamic and predictive biomarker in first-in-human clinical trials.

## Conclusion

The data support repression of immune signalling by cMYC-G9a/EZH2 and show that chemical inhibition of H3K27me/H3K9me maintenance is capable of disrupting cMYC-associated transcriptional networks, and inducing an immune response accompanied by a survival benefit. Such inhibition is a promising anti-tumour approach in *MYC*-deregulated tumours. cMYC status and the 7ISG may predict tumour sensitivity, and support clinical evaluation of inhibition of H3K27 and H3K9 methylation in *MYC*-deregulated tumours.

## Acknowledgements

We would like to thank Dr Edward Curry for initial exploration of the Sanger data and Nahal Masrour for laboratory support. The work was supported by Ovarian Cancer Action, Imperial College Confidence in Concept grant, a Stratigrad Imperial College PhD studentship to ID, Cancer Research UK (A27603) and Medical Research Council (MRC NMGN MC_PC_21042).

