## Supplementary Figures for "cMYC-mediated immune repression is reversed by inhibition of H3K9/H3K27 methylation maintenance"

### Supplementary figure 1

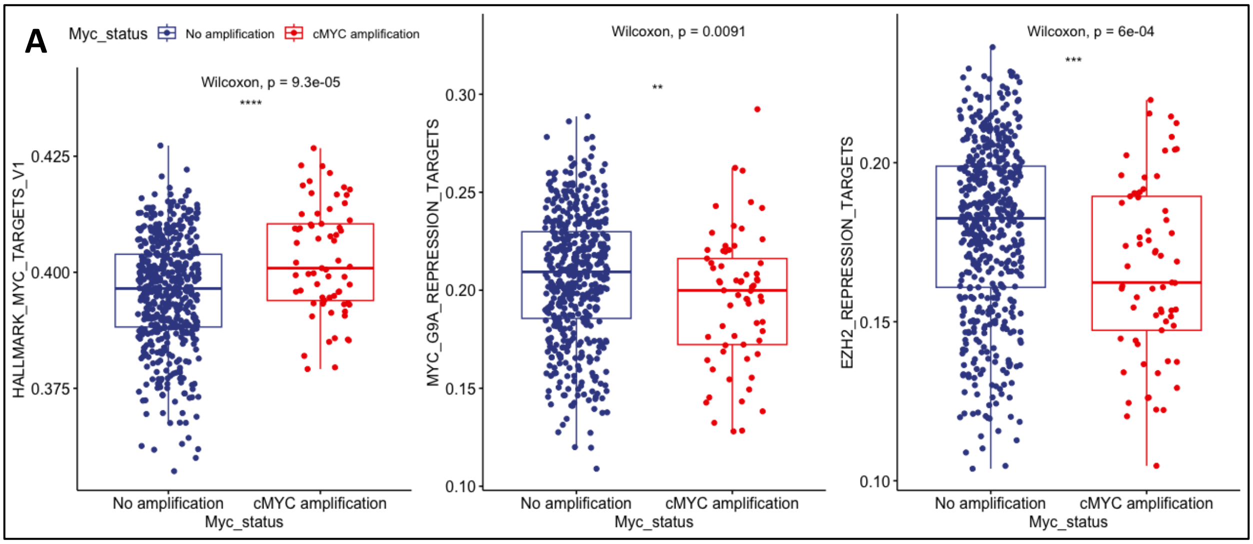


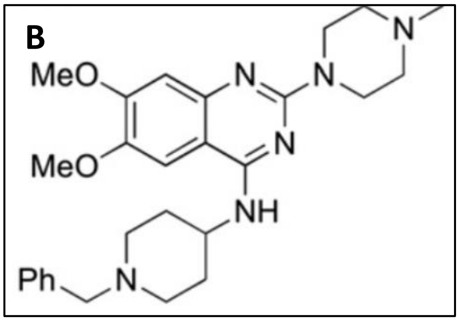

### Supplementary figure 2

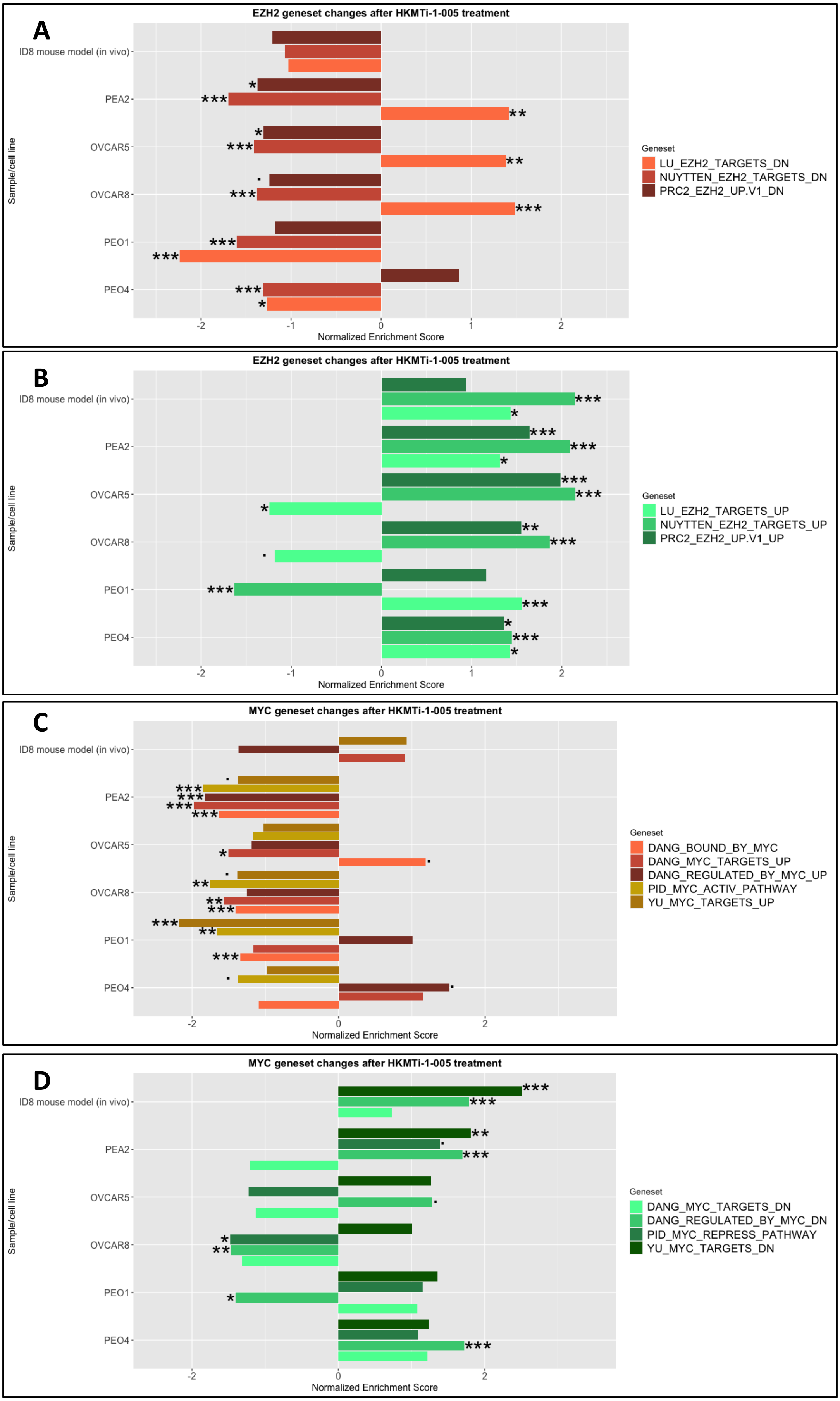

### Supplementary figure 3

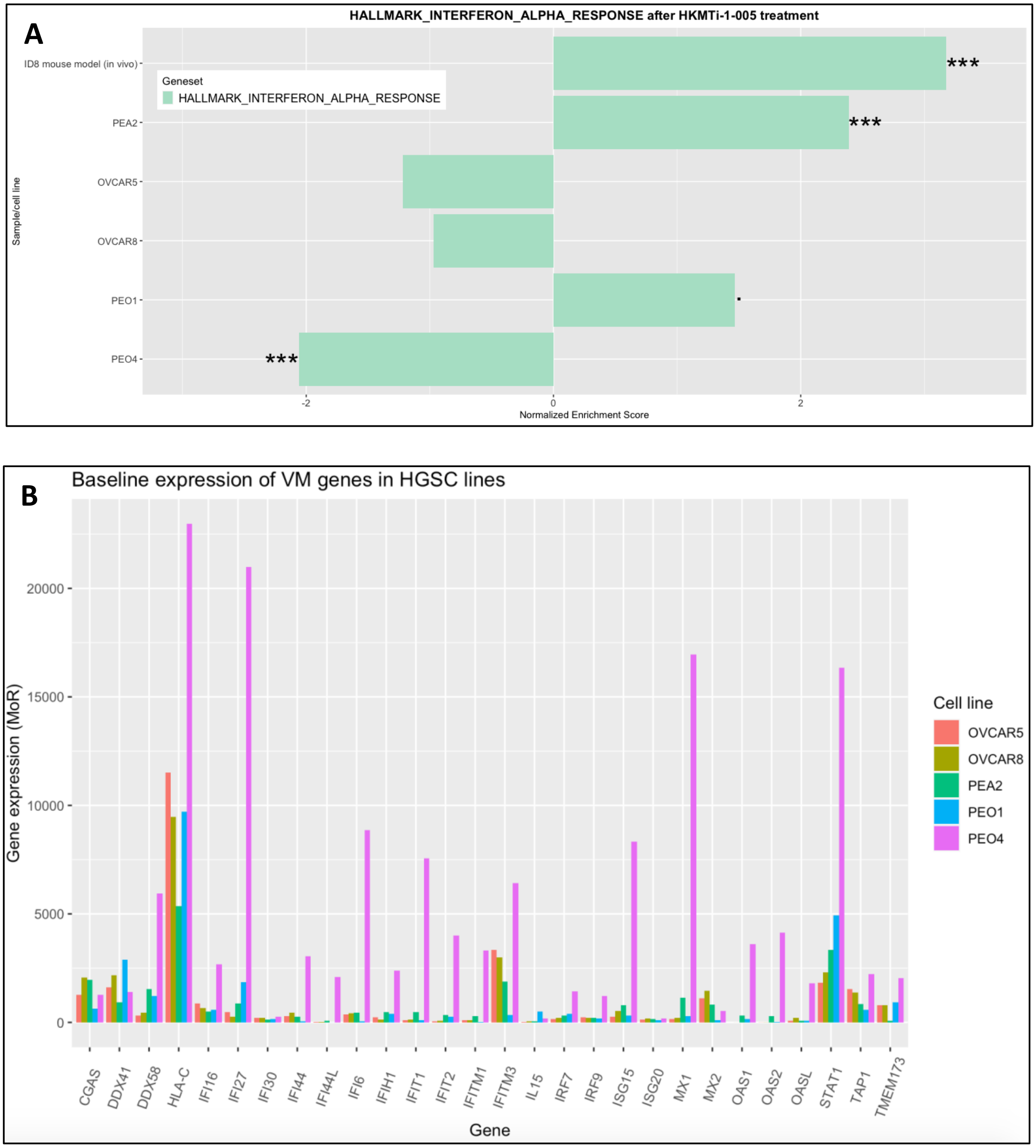

### Supplementary figure 4

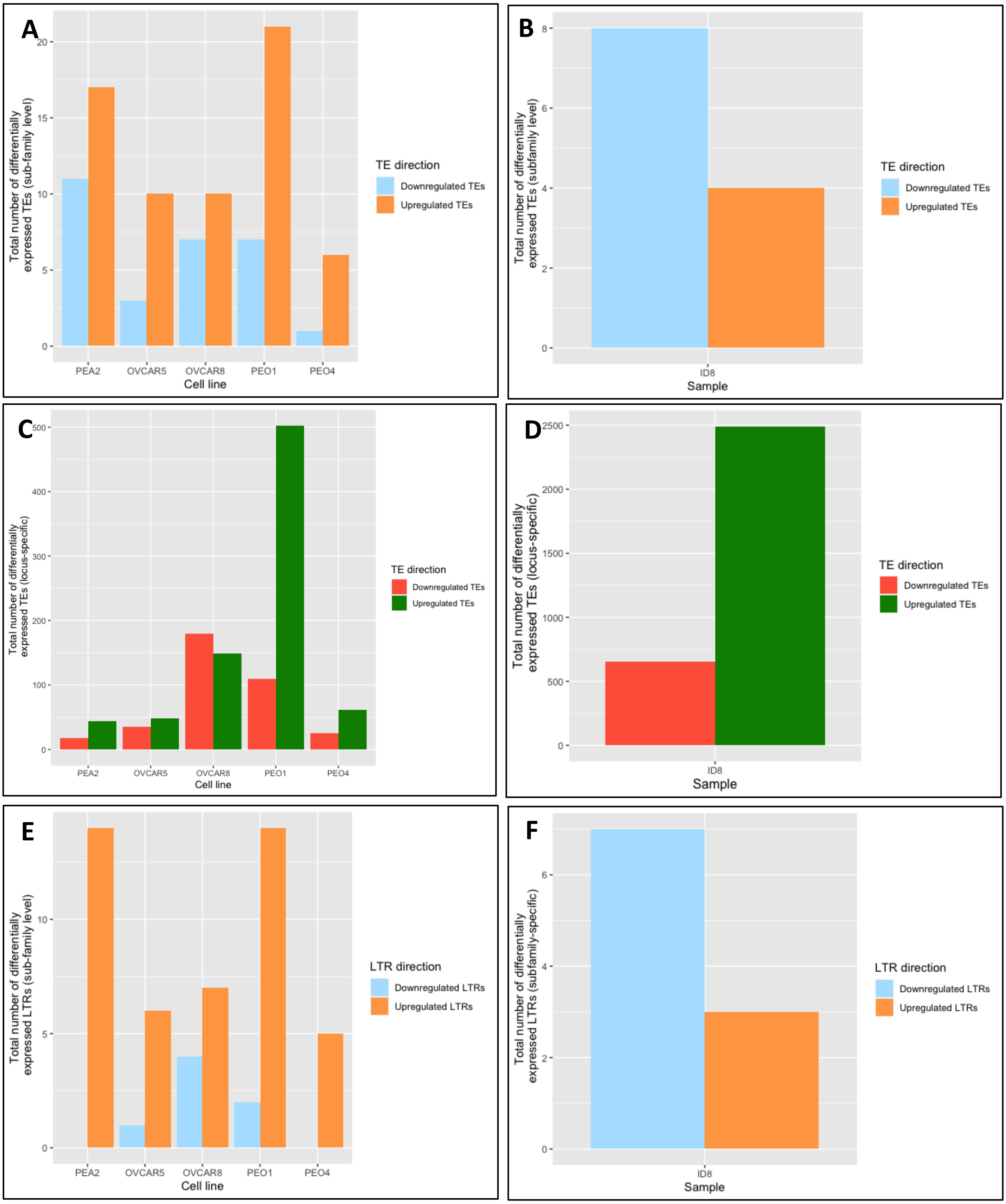

### Supplementary figure 5

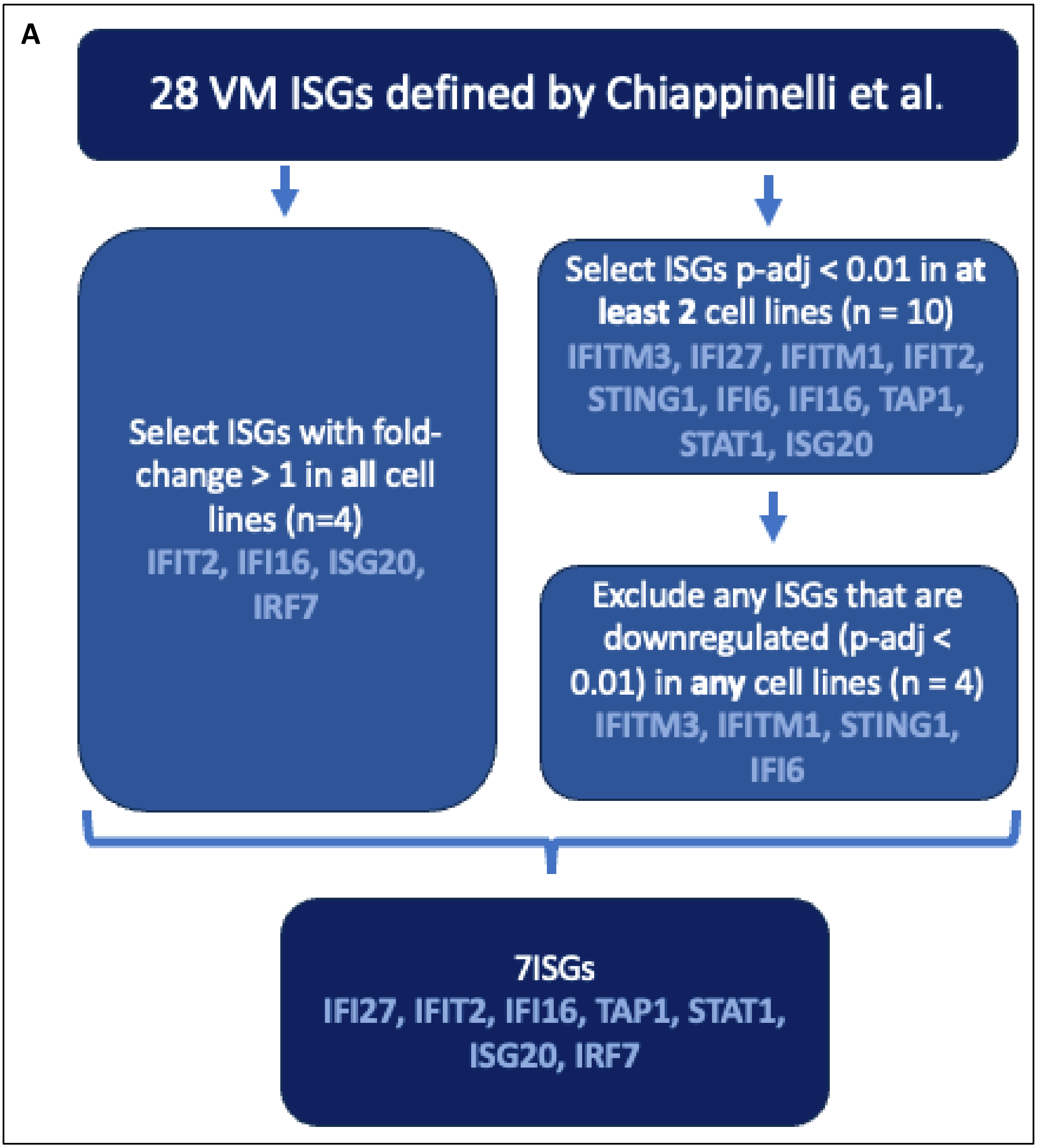

### Supplementary figure 6

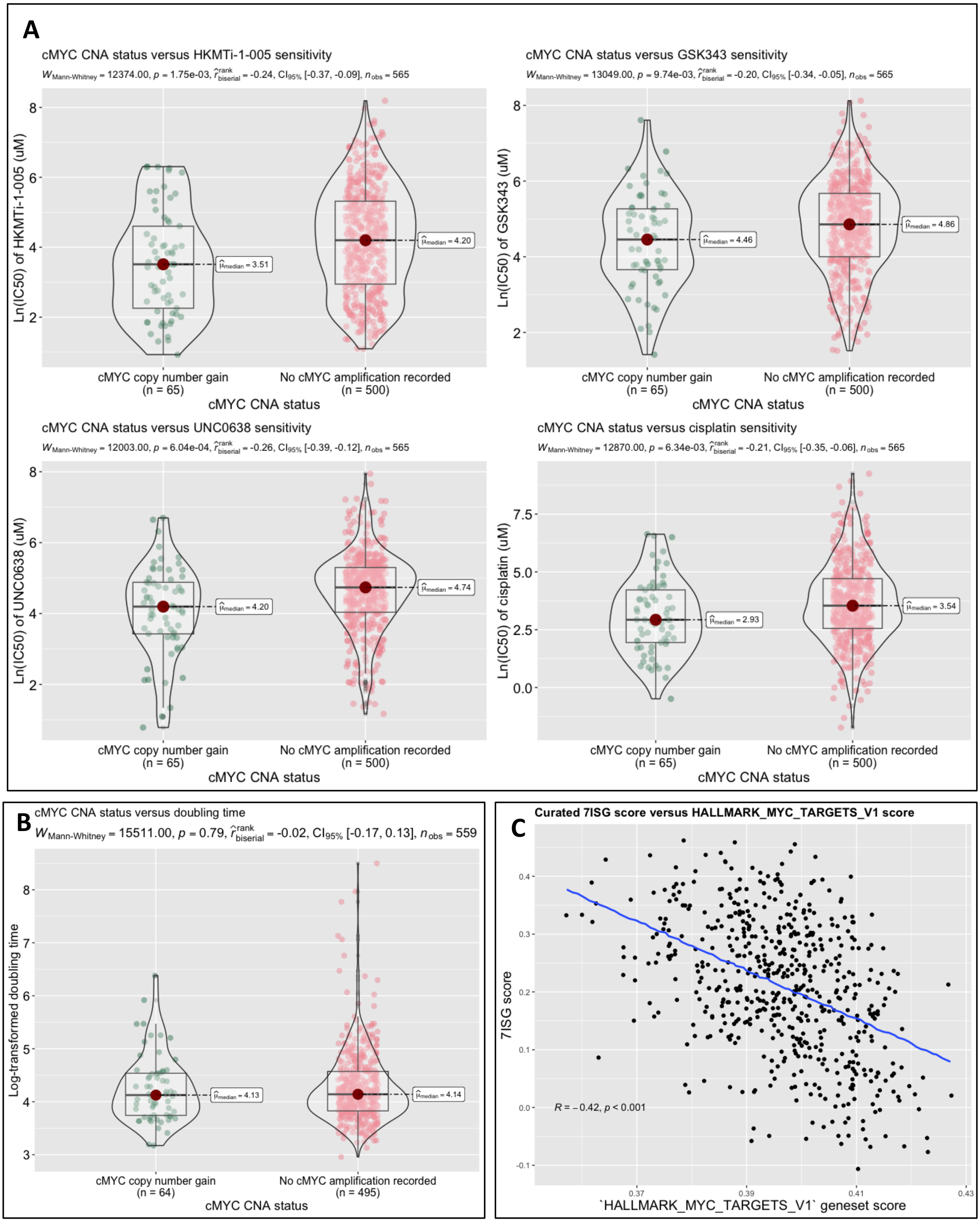

### Supplementary figure 7

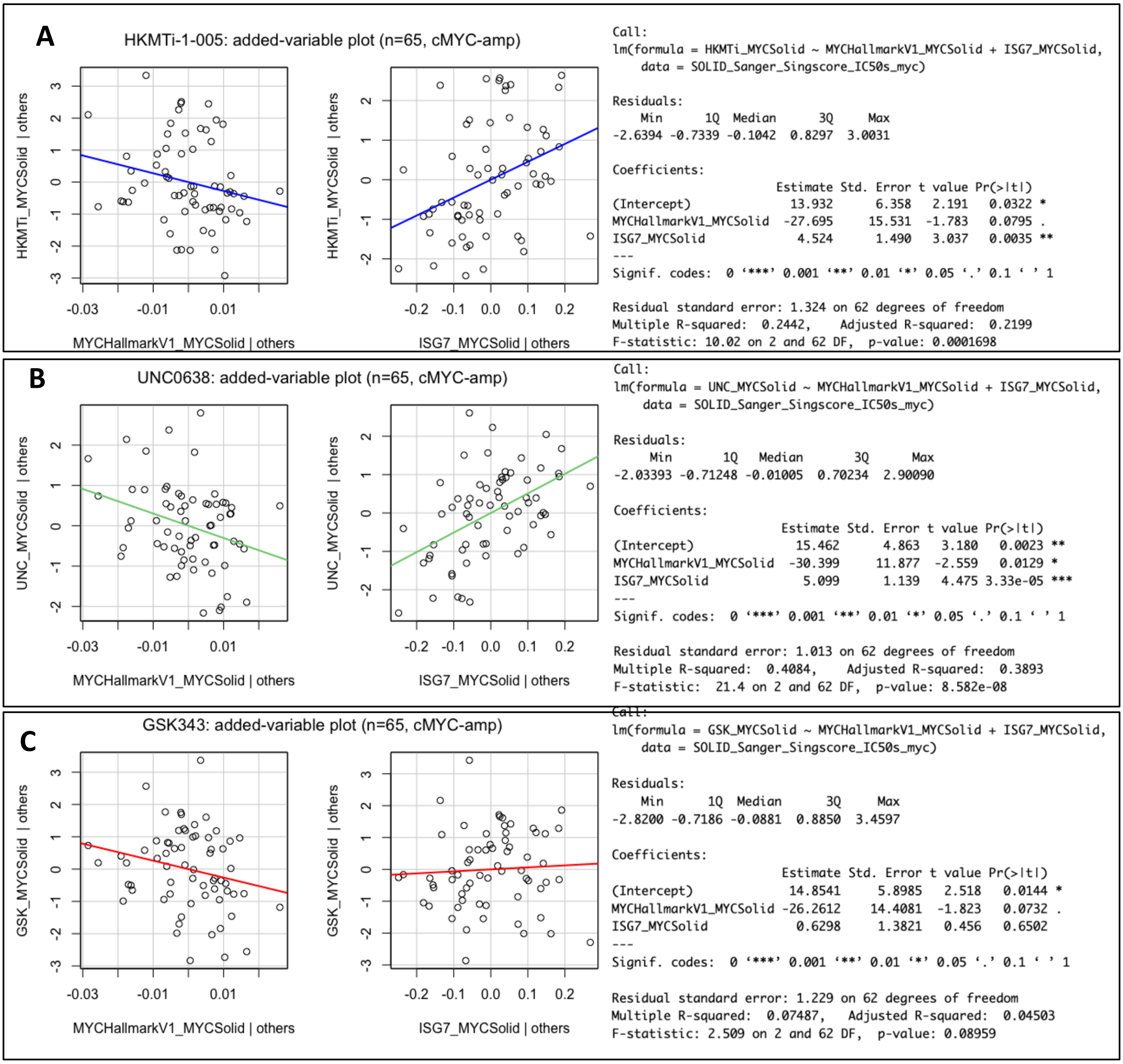

### Supplementary figure 8

A

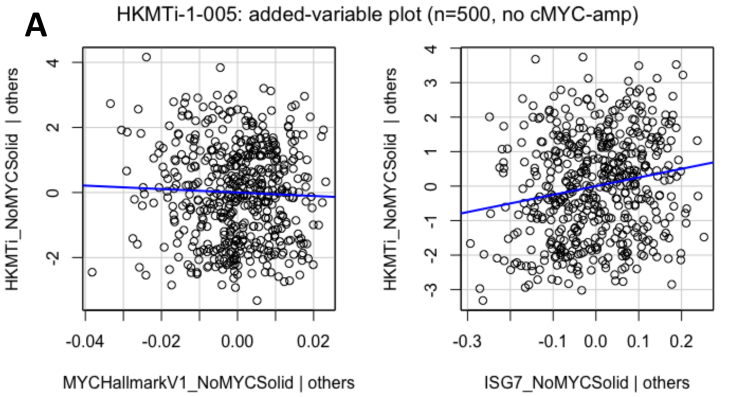


```
Call:
lm(formula = HKMTi_NoMYCSolid ~ MYCHallmarkV1_NoMYCSolid + ISG7_NoMYCSolid,
    data = SOLID_Sanger_Singscore_IC50s_NoMYC)

Residuals:
    Min      1Q  Median      3Q     Max
-3.2938 -1.2299  0.0031  1.1255  4.0345

Coefficients:
                         Estimate Std. Error t value Pr(>|t|)
(Intercept)                5.7278     2.4704   2.319   0.0208 *
MYCHallmarkV1_NoMYCSolid  -5.3046     6.0900  -0.871   0.3842
ISG7_NoMYCSolid            2.5102     0.6249   4.017 6.81e-05 ***
---
Signif. codes:  0 ‘***’ 0.001 ‘**’ 0.01 ‘*’ 0.05 ‘.’ 0.1 ‘ ’ 1

Residual standard error: 1.47 on 497 degrees of freedom
Multiple R-squared:  0.04641,   Adjusted R-squared:  0.04257
F-statistic: 12.09 on 2 and 497 DF,  p-value: 7.432e-06
```

B

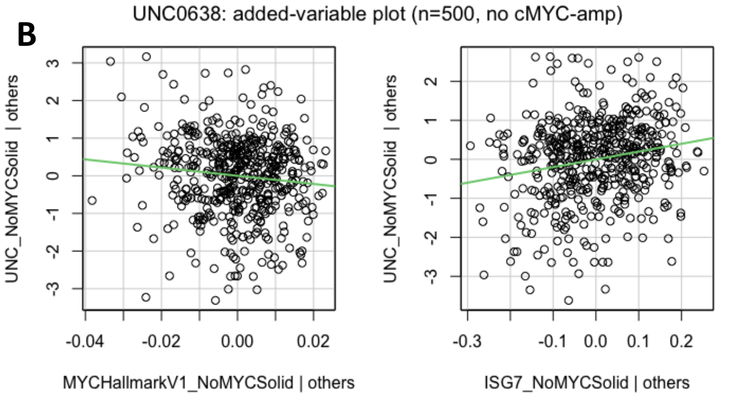


```
Call:
lm(formula = UNC_NoMYCSolid ~ MYCHallmarkV1_NoMYCSolid + ISG7_NoMYCSolid,
    data = SOLID_Sanger_Singscore_IC50s_NoMYC)

Residuals:
    Min      1Q  Median      3Q     Max
-3.4893 -0.6307  0.1059  0.6943  2.9044

Coefficients:
                         Estimate Std. Error t value Pr(>|t|)
(Intercept)                8.4861     1.7627   4.814 1.96e-06 ***
MYCHallmarkV1_NoMYCSolid -10.8647     4.3454  -2.500   0.0127 *
ISG7_NoMYCSolid            1.9975     0.4459   4.480 9.28e-06 ***
---
Signif. codes:  0 ‘***’ 0.001 ‘**’ 0.01 ‘*’ 0.05 ‘.’ 0.1 ‘ ’ 1

Residual standard error: 1.049 on 497 degrees of freedom
Multiple R-squared:  0.08067,   Adjusted R-squared:  0.07697
F-statistic: 21.81 on 2 and 497 DF,  p-value: 8.363e-10
```

C

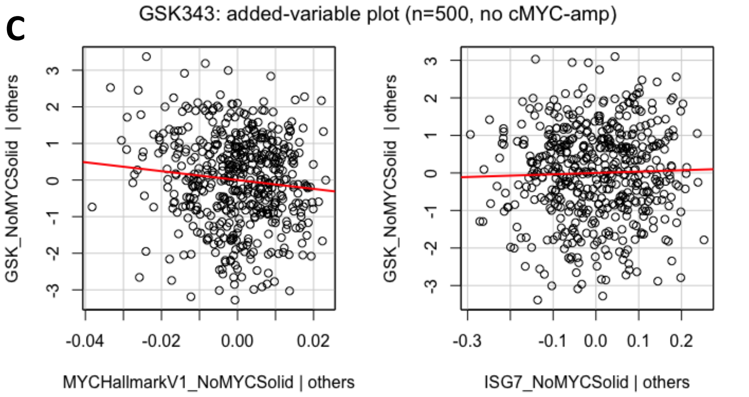


```
Call:
lm(formula = GSK_NoMYCSolid ~ MYCHallmarkV1_NoMYCSolid + ISG7_NoMYCSolid,
    data = SOLID_Sanger_Singscore_IC50s_NoMYC)

Residuals:
    Min      1Q  Median      3Q     Max
-3.3367 -0.7978  0.0456  0.8892  3.0828

Coefficients:
                         Estimate Std. Error t value Pr(>|t|)
(Intercept)                9.5134     2.0702   4.595 5.48e-06 ***
MYCHallmarkV1_NoMYCSolid -12.1153     5.1035  -2.374    0.018 *
ISG7_NoMYCSolid            0.3604     0.5237   0.688    0.492
---
Signif. codes:  0 ‘***’ 0.001 ‘**’ 0.01 ‘*’ 0.05 ‘.’ 0.1 ‘ ’ 1

Residual standard error: 1.232 on 497 degrees of freedom
Multiple R-squared:  0.01803,   Adjusted R-squared:  0.01408
F-statistic: 4.563 on 2 and 497 DF,  p-value: 0.01087
```
